# Pregnancy Reduces Il33+ Hybrid Progenitor Accumulation in the Aged Mammary Gland

**DOI:** 10.1101/2024.08.01.606240

**Authors:** Andrew Olander, Cynthia M Ramirez, Veronica Haro Acosta, Paloma Medina, Sara Kaushik, Vanessa D Jonsson, Shaheen S Sikandar

## Abstract

Aging increases breast cancer risk while an early first pregnancy reduces a woman’s life-long risk. Several studies have explored the effect of either aging or pregnancy on mammary epithelial cells (MECs), but the combined effect of both remains unclear. Here, we interrogate the functional and transcriptomic changes at single cell resolution in the mammary gland of aged nulliparous and parous mice to discover that pregnancy normalizes age-related imbalances in lineage composition, while also inducing a differentiated cell state. Importantly, we uncover a minority population of *Il33*-expressing hybrid MECs with high cellular potency that accumulate in aged nulliparous mice but is significantly reduced in aged parous mice. Functionally, IL33 treatment of basal, but not luminal, epithelial cells from young mice phenocopies aged nulliparous MECs and promotes formation of organoids with *Trp53* knockdown. Collectively, our study demonstrates that pregnancy blocks the age-associated loss of lineage integrity in the basal layer through a decrease in *Il33+* hybrid MECs, potentially contributing to pregnancy-induced breast cancer protection.

## INTRODUCTION

A woman’s risk of breast cancer increases with age (median age at diagnosis = 62 years^1^), but an early first pregnancy (below the age of 30) significantly reduces lifetime risk^2–8^. Previous studies that have examined changes shortly after pregnancy suggest that post-pregnancy mammary epithelial cells (MECs) have a reduced stem cell capacity^9,10^ and increased expression of differentiation markers^11^. On the other hand, the studies that examined changes with aging in nulliparous (no pregnancy) MECs have found an accumulation of dysfunctional luminal progenitors^12^, lineage infidelity^13^, and altered differentiation programs^14,15^. Importantly, there is not a clear understanding of how these processes combine and evolve with aging, i.e., how does pregnancy alter the process of aging in mammary stem/progenitor cells?

Most of the previous studies focused on time points immediately post-pregnancy or 40 days post-involution (∼3.3 human years), yet pregnancy-induced protection does not take effect until almost 10 years post-birth^4,16,17^. To address this gap in knowledge, we simulated conditions that mimic an early first pregnancy (20-30 years in humans, 3-8 months in mice) and the post-menopausal stage (>50 years in humans, ∼18 months in mice). The long-term effect of pregnancy on aging of MECs is important to determine because 75% breast cancer diagnoses occur over the age of 50^18^, while most women in the United States have their first pregnancy between 20 and 33 years of age^19,20^. Using this 18-month time point in conjunction with a 3-month control, we interrogate the aging process of MECs with and without pregnancy while removing confounding factors affecting human samples^21^. Our study not only delineates the combined effects of aging and pregnancy, but also uncovers a previously unknown *Il33+* hybrid MEC population that accumulates with age and is reduced in aged mice that have undergone pregnancy.

## RESULTS

### Pregnancy induces long lasting changes in cell fate decisions and reduces organoid formation

To understand whether pregnancy alters the aging of the mammary gland, we compared 18-month-old parous mice (18m P) that have undergone multiple pregnancies between 3-8 months of age with 3-month-old nulliparous (3m NP) and 18-month-old nulliparous (18m NP) (**Fig. 1a**). H&E staining and whole mount imaging showed no major differences in ductal density, morphology, or complexity of branching between 18m NP and 18m P (**Extended Data Fig. 1a, b**).

**Figure 1.**
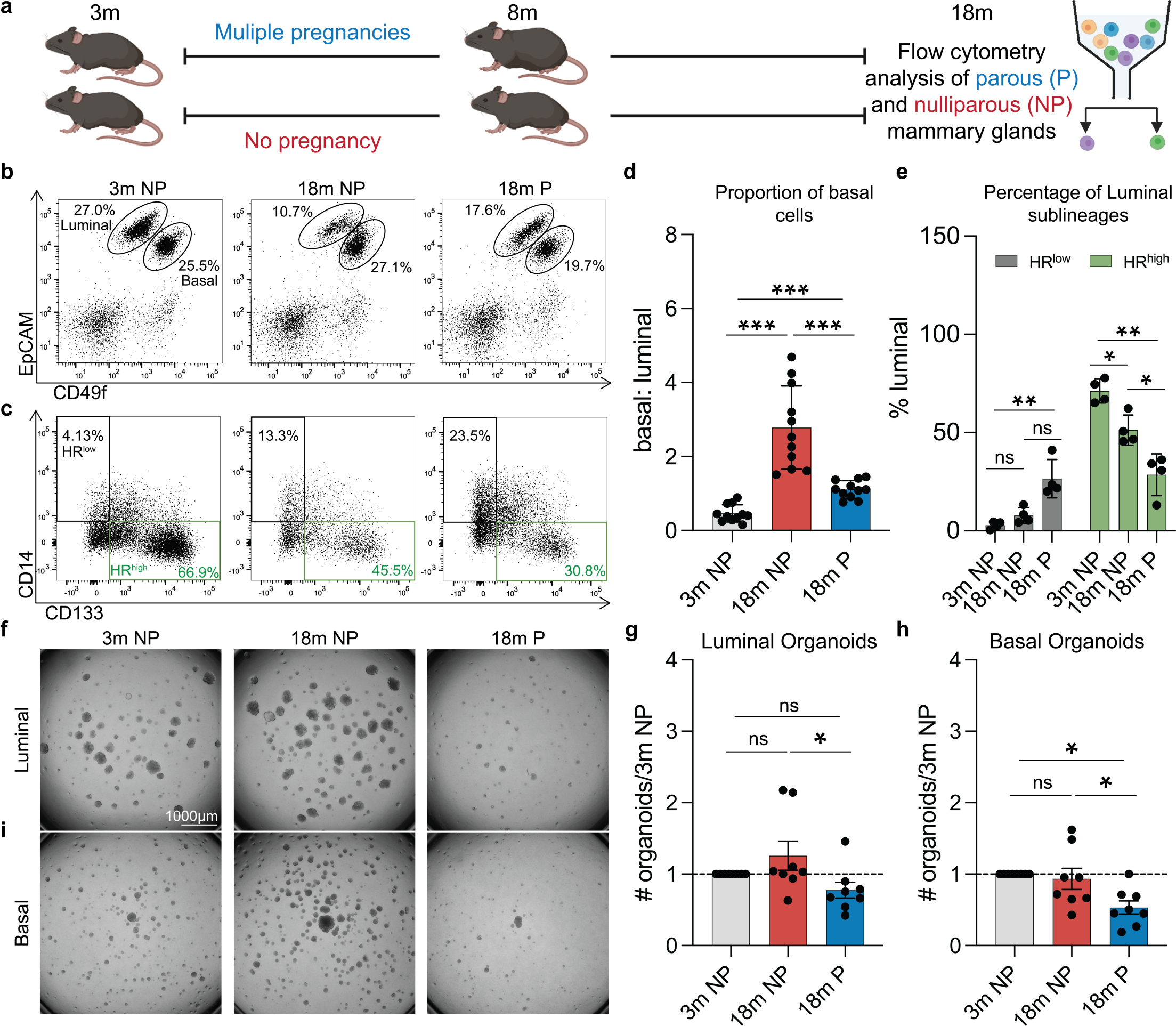
Pregnancy induces persistent changes in cell fate decisions and reduces organoid formation. (a) Schematic of flow cytometry analysis for 3-month nulliparous (3m NP), 18-month nulliparous (18m NP), and 18-month parous (18m P). Created with Biorender.com. (b) Representative flow cytometry plots of DAPI-/CD45-/CD31-/Ter119-single cells from mouse mammary tissues. Basal (EPCAM^med/low^CD49f^high^) and luminal (EPCAM^high^CD49f^med/low^) cells are denoted by representative gates. (c) Quantification of flow cytometry data shown in (b) (n = 11 mice). (d) Representative flow cytometry plots of DAPI-/Lineage-/EPCAM^high^CD49f^med/low^ single cells from mouse mammary tissues. (e) Quantification of flow cytometry data shown in (d) (n = 4 mice). (f) Representative images of primary organoids derived from 3m NP, 18m NP, and 18m P luminal cells. (g) Quantification of luminal organoid number. (h) Quantification of basal organoid number. (i) Representative images of primary organoids derived from 3m NP, 18m NP, and 18m P basal cells. n = 8 mice for experiments in f-i. Statistical significance was determined by performing ANOVA (d, e) or paired t tests (g, i). * p <0.05, ** p < 0.01, *** p < 0.001.

The mammary gland is a bilayered tree-like structure composed of two main epithelial cell lineages, luminal and basal cells^22^. Luminal cells can be further subdivided into two functionally distinct cell types: hormone receptor (HR)-high and -low cells^23^. To understand how these populations evolve with aging and pregnancy, we performed flow cytometry analysis using previously established markers of MEC lineages, basal (CD49F^high^/EPCAM^med-low^) and luminal (CD49F^med^/EPCAM^high^) (**Extended Data Fig. 2**). We found that in 18m NP mice, there was a significant increase in the basal population (increased basal:luminal ratio) as compared to 3m NP mice (**Fig. 1b, d**), consistent with previous studies^24^. However, 18m P mice showed a basal:luminal ratio lower than 18m NP but still higher than 3m NP mice, suggesting that pregnancy partially normalizes the aged-induced expansion of basal cells (**Fig. 1b, d**).

Within the luminal compartment, our analysis showed a minor but non-significant increase in HR-low luminal cells (CD14+/CD33-) and a significant decrease in HR-high luminal cells (CD14-/CD133+) in the 18m NP mice (**Fig. 1c, e**), consistent with previous studies in 13-14 month mice^23^. However, in contrast to the normalization of basal:luminal ratio in 18m P mice, we found a further increase in HR-low and a decrease in HR-high luminal cells in these mice (**Fig 1c, e**). Thus, our data suggests that while the age-induced expansion of basal cells is normalized by pregnancy, the luminal cells retain a residual involution program, with an increased proportion of CD14+ HR-low luminal cells.

To understand whether there are functional differences in luminal and basal cells between the 3m NP, 18m NP and 18m P mice, we performed *in vitro* organoid assays. In contrast to previous studies that have shown an age-induced increase in organoid formation^24^, we found that 3m and 18m NP MECs had similar capacity to form organoids (**Fig. 1, f-i**). However, both luminal and basal cells from 18m P mice had a significantly decreased capacity of organoid formation as compared to 18m NP mice (**Fig. 1f-i**). These findings suggest that MECs from the aged parous glands are diminished in clonogenicity, supporting the hypothesis that pregnancy reduces regenerative potential.

### Pregnancy promotes a differentiated cell state but also reverses aged-associated transcriptional programs

To determine transcriptomic changes in response to aging and pregnancy, we sorted luminal and basal cells and performed bulk RNA sequencing from 3m NP, 18m NP, and 18m P mice. Principal component analysis (PCA) of gene expression values showed that luminal and basal samples formed two distinct clusters (**Fig. 2a**), as expected. Moreover, gene expression for established lineage markers showed that sorting was specific for basal and luminal cells, with enrichment of *Krt14*, *Krt5*, and *Acta2* in the basal cells and *Krt8*, and *Krt18* in luminal cells (**Fig. 2b**).

**Figure 2.**
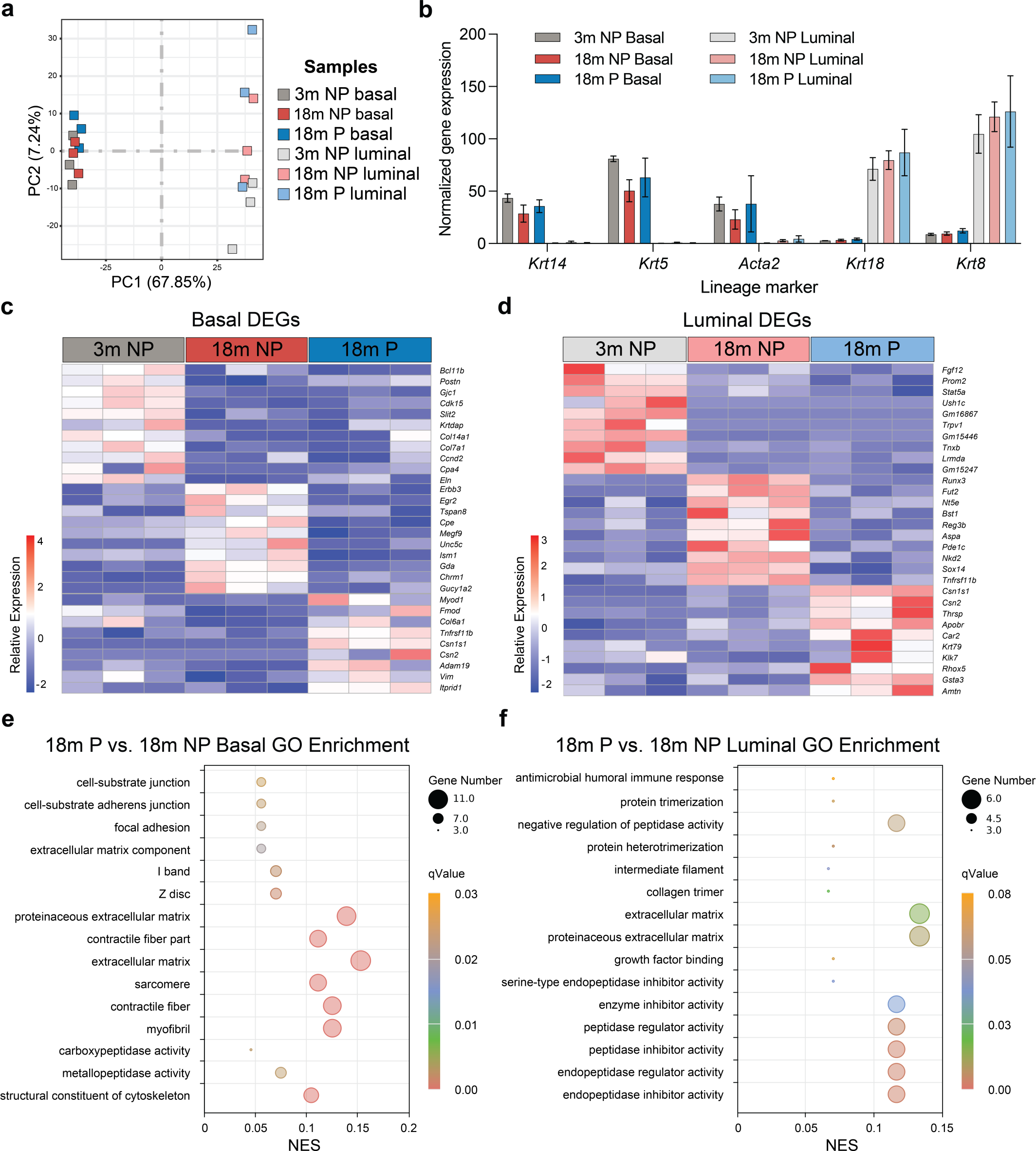
Bulk RNA sequencing in aged nulliparous and parous mammary glands reveals long-lasting transcriptomic changes. (a) Principal Component Analysis (PCA) of basal and luminal RNA-seq samples from 3m NP, 18m NP, and 18m P mice. (b) Bar plot depicting the expression (FPKM x10^3^) of basal (*Krt14, Krt15, Acta2*) and luminal (*Krt18, Krt8*) marker genes across samples. (c) Heatmap of differentially expressed genes (DEGs) across 3m NP, 18m NP, and 18m P basal cells and (d) luminal cells. Relative expression reflects log2(FC+1) values. Significance was determined using an adj-p value of <= 0.1. (e, f) Dotplot of Gene Ontology (GO) results for 18m P basal (e) and luminal (f) cells, relative to 18m NP. GO results were selected from the top 15 hits with the lowest adj-p value denoted as q-value (< 0.03 for basal, < 0.08 for luminal), NES = Normalized Enrichment Score. Gene number refers to the number of genes from the input gene list that are present in the GO gene list.

To understand the long-term effects of pregnancy on gene expression in MECs, we performed differential gene expression analysis between 18m NP and 18m P for basal and luminal cells. Although luminal cells undergo significant functional and transcriptomic changes throughout pregnancy^25^, we found only 45 differentially expressed genes (DEGs) at p-adj of =< 0.1 that are persistently altered in luminal cells (**Data Table 1**). Interestingly, we found 74 DEGs in basal cells at the same p-adj threshold (**Data Table 2**). Previous studies suggest that parity has the most pronounced impact on the transcriptome of luminal cells^25^, but our data suggest that basal cells also retain a robust transcriptional memory of pregnancy that persists with age. Basal cells from 18m P mice displayed consistent upregulation of genes associated with differentiation towards a mesenchymal lineage and contractile functions, such as *Myod1, Fmod,* and *Vim* (**Fig. 2c**). Moreover, 18m P basal cells had increased expression of milk-protein genes, such as *Csn1s1* and *Csn2*, suggesting that basal cells upregulate pathways of milk-production during pregnancy or give rise to milk producing cells^26^, or vice versa^27^. We also detected a modest down regulation of genes associated with stem/progenitor cells (*Tspan8* and *Bcl11b*)^28,29^ and growth factor receptors (*Erbb3* and *Egr2*). Likewise, luminal cells from 18m P mice had higher expression of genes involved in alveolar differentiation (*Csn1s1, Csn2, Car2,* and *Apobr*) and down regulation of genes involved in stem/progenitor function (*Runx3* and *Bst1*, **Fig. 2d**). Taken together, these results suggest that pregnancy establishes long-term transcriptomic changes that reflect increased differentiation not only in luminal cells but also in basal cells. Interestingly, by comparing the gene expression patterns between 3m NP, 18m NP and 18m P mice we found that expression of *Tspan8*, that marks hormone-responsive stem cells^28^ and growth factor receptors (*Erbb3* and *Egr2*) increases with aging in basal cells but is reduced in 18m P mice (**Fig. 2c**). Similar patterns were observed for luminal cells in stem/progenitor genes (*Runx3* and *Bst1*), the RANKL decoy receptor, *Tnfrsf11b,* and the *Wnt* inhibitor, *Nkd2*, among others (**Fig. 2d, see** Data tables 1-4).

To identify specific pathways that are altered by pregnancy and persist with age, we performed gene ontology analysis on basal or luminal cells in 18m P and 18m NP mice. We found significant enrichment of genes involved in cytoskeletal remodeling, extracellular matrix, and contractile functions (**Fig. 2e**). Similarly, 18m P luminal cells showed enrichment of pathways involved in regulation of peptide metabolism, antimicrobial immune responses, and collagen structure (**Fig. 2f**), which have been proposed to play key roles during alveologenesis, involution^30^, and immune development of offspring^31,32^. These data suggest that pregnancy normalizes age-induced transcriptional changes in luminal and basal epithelial cells, while also inducing a differentiated state.

### Single-cell RNA-sequencing identifies a minority population of hybrid MECs in aged nulliparous mice

Recent studies have demonstrated that minority populations of stem/progenitor cells evolve with age and are likely precursors of tumor initiation^24,33^. To investigate the cellular heterogeneity and aging-associated transcriptional changes that are altered with pregnancy at single-cell resolution, we performed single-cell RNA-sequencing (scRNA-seq) on mammary glands from 18m NP and 18m P mice. To increase the power of our analysis, we integrated single cell transcriptomes from the Tabula Muris^34^ 18m mammary gland dataset (see methods). After filtering cells with low-expressing genes, high mitochondrial counts, and doublets (see methods), we identified epithelial cells (basal, HR-high luminal, HR-low luminal), immune cells (B cells, T cells, macrophages), endothelial cells, and fibroblasts using the expression of well-established marker genes (**Fig. 3a, Extended Data Fig. 3a**). Intriguingly, we identified a minority population of cells co-expressing basal and luminal markers (hybrid) (**Fig. 3c**), that are present in 18m NP mice (∼90% of cell cluster) but are significantly reduced in 18m P mice (∼10% of cell cluster) (**Fig. 3b**). Moreover, we found no differences in the proportion of B cells, T cells, macrophages, endothelial cells, and HR-low luminal cells between 18m P and 18m NP (**Fig. 3b**). However, 18m P had a lower proportion of basal cells and HR-high luminal cells (**Fig. 3b**), consistent with our flow cytometry analyses (**Fig. 1b-e**). Moreover, we did not find differences in cell cycle between 18m NP and 18m P mice (**Extended Data Fig. 3b**).

**Figure 3.**
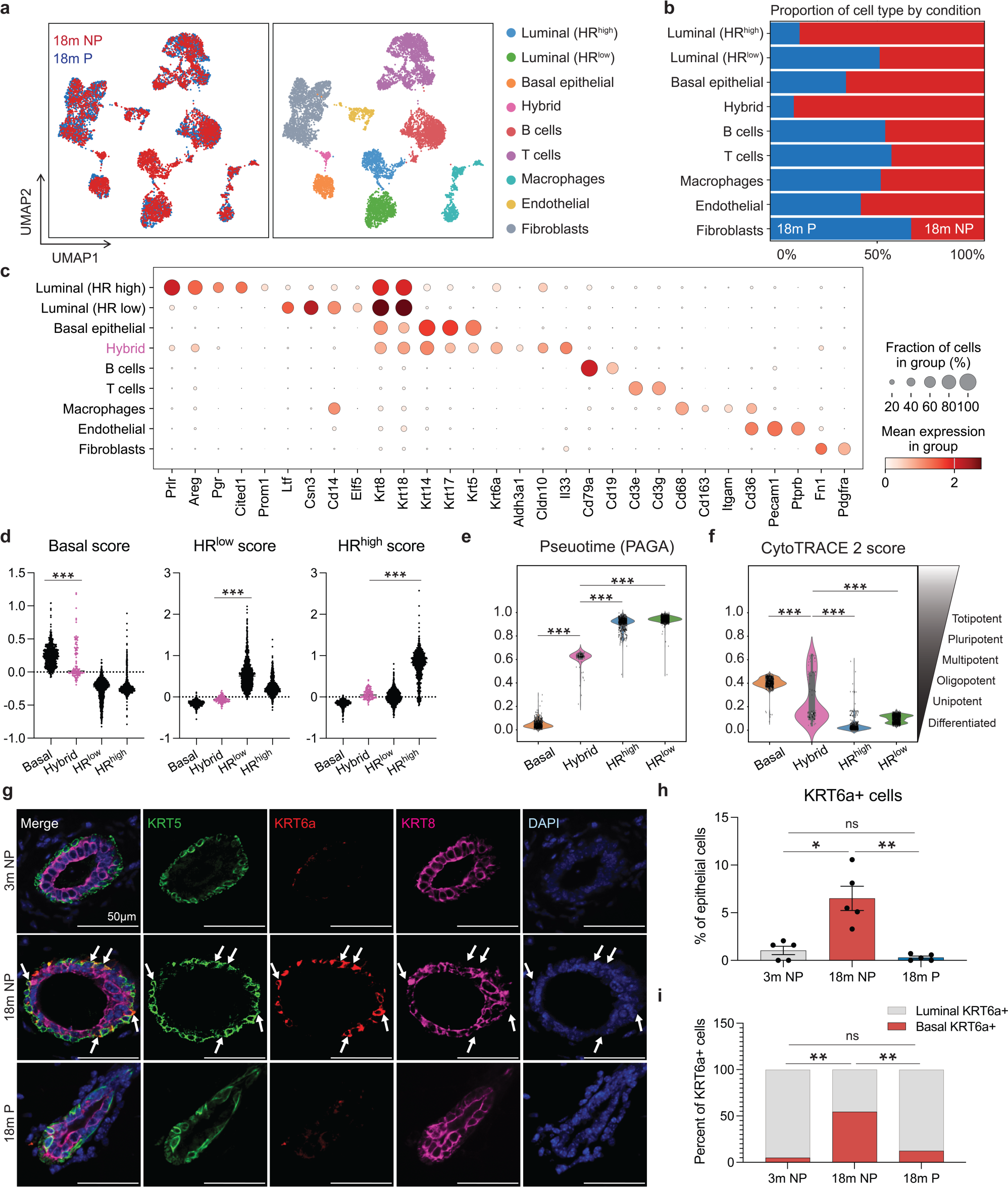
Single cell RNA sequencing identifies a hybrid population of MECs in aged nulliparous mice but is reduced in aged parous mice. (a) UMAP plot of scRNA-seq data generated from the mammary glands of 18m nulliparous (NP, red) and parous (P, blue) mice (n = 3 mice) and integrated with Tabula Muris (n=10,001). UMAPs are colored by condition (left) and annotated cell type (right). (b) Bar plot showing the proportion of cell types by condition. (c) Dotplot of gene expression for selected marker genes of mammary cell lineages. (d) Basal, HR-low, and HR-high gene expression scores across epithelial clusters (black), including a minority population of hybrid MECs (pink). (e) Violin plot showing the distribution of inferred partition-based graph abstraction (PAGA) pseudotime scores across epithelial cell clusters. (f) Violin plot comparing predicted developmental potential (obtained through CytoTRACE 2) across epithelial cell clusters. (g) Representative immunofluorescence (IF) stains against KRT5 (basal cells, green), KRT6a (hybrid MECs, red), and KRT8 (luminal cells, magenta). Nuclei are visualized using DAPI (blue). Hybrid MECs localized to the basal layer are indicated by white arrows. (h) Quantification of all KRT6a+ hybrid MECs in the epithelium. (i) Quantification of the localization of KRT6a+ hybrid MECs (basal or luminal). Statistical significance was determined by performing one-way ANOVA with Tukey test (d) or unpaired t tests with Welch’s correction to account for unequal standard deviations (h). * p <0.05, ** p < 0.01, *** p < 0.001. Statistical tests for scRNA-sequencing are described in the methods section. n = 5 mice for IF experiments. Scale bar = 50 µm.

To investigate the unknown hybrid population further, we calculated basal cell and luminal cell gene signature enrichment scores for all cells based on previously published transcriptomic analyses of adult MECs^35^. We found that the hybrid MEC population lies between the luminal and basal populations, implying that hybrid MECs display a mixed gene signature of both basal and luminal lineages (**Fig. 3d**). Pseudotime inference using PAGA^36^ and Slingshot^37^ also predicted the hybrid MECs lineage trajectory between basal and luminal populations (**Fig. 3e, Extended Data Fig. 4a, b**).

To understand whether these hybrid MECs represent an immature cell type, we applied CytoTRACE2^38^, to agnostically predict cellular potency. Intriguingly, we found that 33% hybrid MECs are predicted to be multipotent, while majority of basal and luminal cells were predicted to be oligopotent and differentiated, respectively. Moreover, the average cellular potency of hybrid MECs was between basal and luminal cells (**Fig. 3f, Extended Data Fig. 4c**). Intriguingly, the hybrid MECs are also enriched in expression of *Krt6a* (**Extended Data Fig. 5a**), which has been previously identified as a marker of bipotent luminal progenitor cells in the mammary gland^39^. KRT6a+ cells are normally found only in the luminal layer and do not express the basal marker *Krt14*. However, a previous report showed that a population of KRT14+/KRT6a+ cells expands during early stages of pregnancy^40^. Analysis of the Pal et al. (2017) scRNA-seq dataset containing all mammary gland developmental stages showed that *Krt6a* expression peaks during early pubertal development, particularly within terminal end buds (TEBs)^41^ (**Extended Data Fig. 5b, c**). Taken together, these data suggest that aging coincides with aberrant activation of *Krt6a* expression likely conferring a cell state similar to TEBs, but this is diminished by pregnancy.

### Hybrid KRT6a+ cells are present in the basal epithelium of aged nulliparous mice but not aged parous mice

To expand our analysis from mRNA to protein, we performed immunofluorescence staining on 3m NP, 18m NP, and 18m P mammary glands for KRT6a, together with KRT8 (luminal marker) and KRT5 (basal marker). We found a greater than 6-fold increase in the percentage of KRT6a+ cells in the 18m NP (6.51%) mice relative to 3m NP (∼1.05%) (**Fig. 3g, h**), suggesting that these hybrid MECs accumulate over the course of aging. However, 18m P mice had a small percentage of KRT6a+ cells (<1%), less than the 3m NP (**Fig. 3g, h**), consistent with our single-cell analysis. Moreover, we found an increased percentage of KRT5+ basal cells and decreased percentage of KRT8+ luminal cells in 18m NP mice (**Extended Data Fig. 5d**) consistent with our flow cytometry data (**Fig. 1b, d**), but 18m P cells were more similar to 3m NP cells.

To further understand the localization of the KRT6a+ cells, we quantified the percentage of KRT6a+ cells in the luminal and basal layer (i.e., KRT6a+/KRT8+ or KRT6a+/KRT5+). We found that 95% of KRT6a+ cells in 3m NP glands were positive for KRT8 and resided in the luminal layer (**Fig. 3i**), consistent with previous studies^39^. In contrast, over 50% of KRT6a+ cells in the 18m NP gland were positive for KRT5 and localized to the basal layer. Interestingly, in the 18m P mice, the localization of the KRT6a+ cells was closer to 3m NP mice (∼10%, **Fig. 3i**). This suggests that the KRT6a+ hybrid MECs that accumulate with age are likely due to the loss of lineage integrity of basal cells, but this integrity is maintained in mice that have undergone pregnancy.

### IL33 treatment of young basal cells phenocopies aged nulliparous mammary glands

In addition to *Krt6a*, the hybrid MEC population identified in 18m NP mice was uniquely enriched in the expression of *Interleukin 33 (Il33)* (**Fig. 3c, Extended Data Fig. 6a**), also known as alarmin. IL33 has been previously implicated in activating pro-tumorigenic signaling pathways^42–44^, cancer stem cell maintenance^45^, and establishing an immunosuppressive microenvironment^46^, but its role in the normal mammary gland is unknown. Analysis of the Pal et al. (2017) scRNA-seq dataset showed that *Il33* is also highly expressed in the TEBs during puberty, (**Extended Data Fig. 6b, c**), similar to Krt6a (**Extended Data Fig. 5b, c**) suggesting that aging likely upregulates an early developmental program in basal cells.

To determine whether IL33 treatment affects primary MECs *in vitro*, we cultured basal (CD49f^hi^/EPCAM^lo/med^) or luminal (CD49f^lo^/EPCAM^hi^) cells from 3m NP mice with recombinant IL33. Treatment of basal and luminal cells with IL33 resulted in an increased number of basal-derived organoids (**Fig. 4a**) but no increase in organoids derived from luminal cells (**Extended Data Fig. 6d**). Interestingly, IL33 treatment of basal organoids for 2 weeks inhibited spontaneous *in vitro* luminal cell differentiation, resulting in a ∼2-fold increase in the proportion of basal cells (**Fig. 4b**). Moreover, IF staining and 3D imaging demonstrated an increase in the percentage of KRT6a and KRT5 upon treatment with IL33 (**Fig. 4c, Extended Data Fig. 6e**), but no significant change in KRT8+ luminal cells (**Extended Data Fig. 6f, g**). These results suggest IL33 treatment of 3m NP basal cells phenocopies the increased proportion of basal cells and accumulation of KRT6a+ hybrid MECs observed in aged nulliparous mice (**Fig. 3i**).

**Figure 4.**
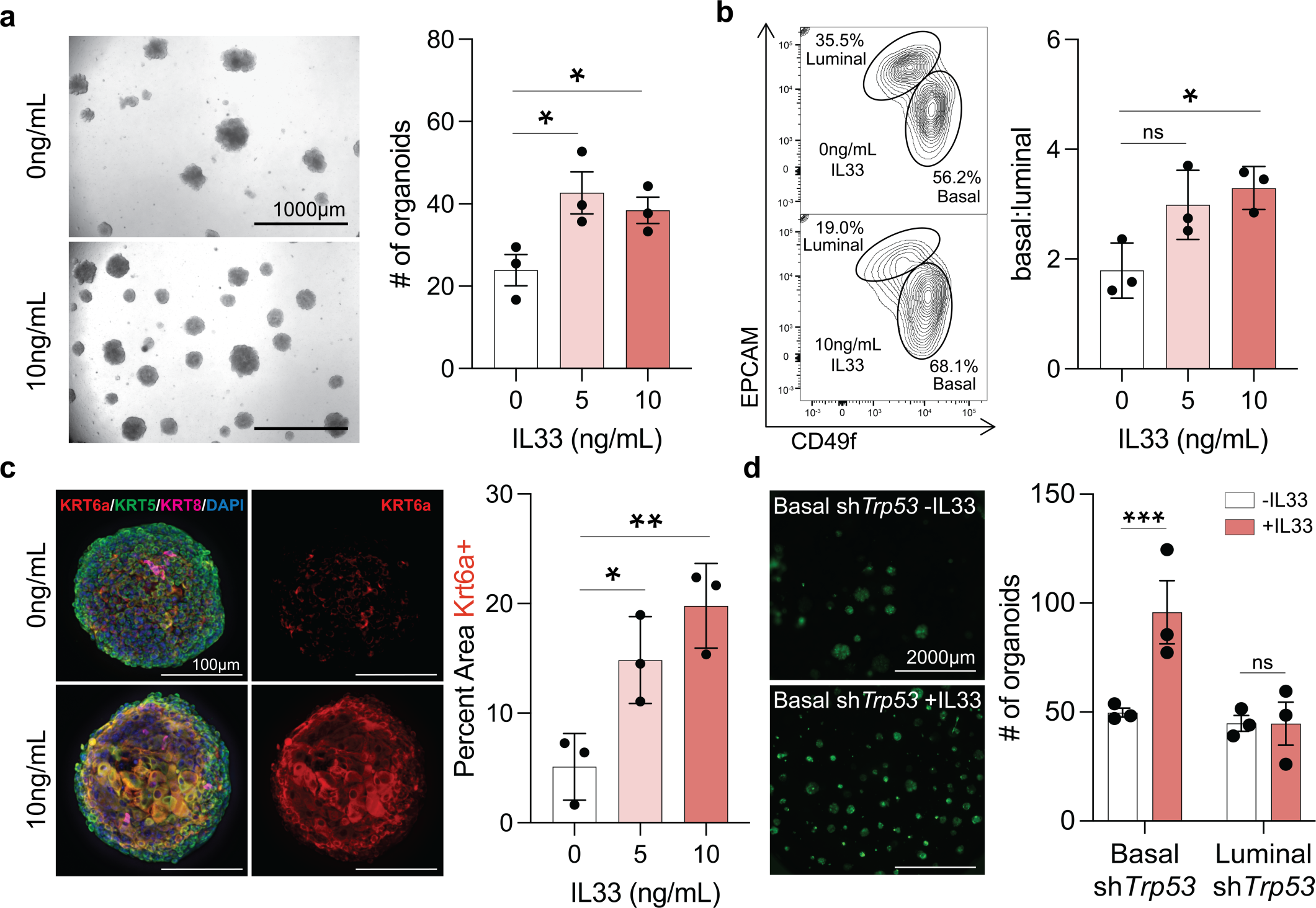
IL33 treatment of young basal cells phenocopies aged nulliparous MECs. (a) Representative images of primary basal organoids from 3m NP mice treated with no Il33 treatment (top left) and with IL33 treatment (bottom left). Quantification of the number of organoids formed with 0, 5, or 10ng/mL of IL33. (b) Representative flow cytometry plots of basal and luminal cells from IL33-treated organoids (left) and the quantified proportion of basal cells relative to the luminal population (right). (c) Representative IF stains (left) against KRT5 (basal cells), KRT6a (hybrid MECs), and KRT8 (luminal cells) on IL33-treated organoids. Nuclei are visualized using DAPI. Quantification of KRT6a+ area in basal organoids cultured with 0, 5, or 10ng/mL of IL33 (right). (d) Representative images of preneoplastic (sh*Tp53*) basal organoids treated with (top left, 10ng/mL) or without IL33 (bottom left). Quantification of the number of sh*Trp53-* basal organoids formed with or without IL33 treatment after 12 days in culture (right). Statistical significance was determined by performing unpaired t tests (a-c) or 2-way ANOVA with Šídák’s multiple comparisons test. * p <0.05, ** p < 0.01, *** p < 0.001. n = 3 mice. Scale bar = 1000 µm (a, d) and 100 µm (c).

To test whether IL33 treatment increases clonogenicity in cells with tumor-promoting mutations, we infected 3m NP basal and luminal cells with lentiviral shRNA targeting *Trp53*^47^ and exposed them to IL33. Consistent with our results in normal epithelial cells (**Fig. 4a, Extended Data Fig. 6d**), we found that *Trp53* knockdown resulted in increased clonogenicity of basal cells when treated with IL33 but not luminal cells (**Fig. 4d**). Thus, our data suggest that the age-induced increase in *Il33*+ hybrid MECs likely confers a survival advantage as cells acquire oncogenic mutations.

## DISCUSSION

Our study resolves the complex relationship between aging and pregnancy in the mammary gland, such as basal-cell bias and differentiation. We report that pregnancy not only normalizes the age-induced expansion of basal cells but also concurrently reduces the capacity of basal cells to form organoids. In contrast, luminal cells in aged parous mice retain an involution specific signature, which potentially makes them more susceptible to immune surveillance^48–51^.

Hybrid MECs expressing luminal and basal genes have been identified previously and are associated with a less-differentiated phenotype, higher plasticity, and are capable of tumor initiation^52–55^ but precise molecular signatures of these cells are not yet defined. Our work identifies a previously unknown *Il33+* hybrid MEC population that accumulates with age in the basal layer. Majority of studies on breast tumor initiation implicate luminal cells as the cell of origin^56^, that lose lineage integrity to express basal markers during tumor initiation and also with aging^12–15^. However, our study suggests that basal cells can acquire hybrid features with aging by turning on pathways active in luminal cells during early pubertal development. In support of this model, studies have shown that basal cells can acquire hybrid phenotypes upon transformation^57^ and form luminal-like tumors^52,54^, which are the predominant subtypes found in postmenopausal breast cancers^58^. Future work will determine the precise contribution of basal cells to tumor initiation with aging.

The postnatal mammary gland development and maintenance is largely driven by lineage committed progenitor cells that expand during pregnancy to form the milk-producing cells^59–61^. Studies have also shown that there are rare bipotent mammary stem cells in the basal layer that can expand with pregnancy and contribute to the alveolar lineage^61,62^. We speculate that during pregnancy some basal cells either turn on an alveolar gene expression program or directly contribute to the alveolar lineage, leading to the normalization of the basal:luminal ratio and a decrease in the hybrid MEC population in aged parous mice.

IL33 has been extensively studied in inflammation and immune cell responses^63^ but its role in epithelial cell plasticity is unknown. Our results show that IL33 increases organoid formation capacity and induces a KRT6a+ hybrid state in basal cells. Recent work has uncovered a role for IL33 in pancreatic cancer where *Il33* expression is induced post-injury and cooperates with mutant *Kras* to promote neoplastic transformation^64^. Notably, when cells acquire hybrid signatures due to expression of *Pik3ca^H1047R^*in luminal cells, they dramatically increase the expression of *Il33*^54^. Moreover, studies have demonstrated a cell-intrinsic role of IL33 in B-cell development^65^, repair during skin injury^66^, and promoting inflammation in chronic pancreatitis^67^. These studies, combined with our data, suggest that *Il33* expression promotes a cell-state that is proliferative and plastic, allowing for injury repair or reprogramming during transformation. Future studies will determine the cell intrinsic role of *Il33* during mammary gland development, in establishing a hybrid cell state, and tumorigenesis.

## METHODS

### Mice

Wild-type C57BL/6 mice (young), retired breeders and aged matched mice when appropriate were purchased from Charles River Laboratories and The Jackson Laboratory. Simultaneously an aging colony was maintained in house. The mice were recorded for estrous and 18m mice were found to be non-cycling. All mice used for this study were maintained at the UCSC Animal Facility/Vivarium in accordance with the guidelines of the Institutional Animal Care and Use Committee (Protocol #Sikas2311dn).

### H&E staining and quantification

H&E staining was performed as previously described^68^. Briefly, mammary glands were dissected from parous and nulliparous mice, fixed in 4% PFA and paraffin embedded for histology. Sections underwent a standard staining protocol, with five-minute incubation in 100% xylene and ethanol (100%, 70%) followed by DI water for deparaffinization. Sections were stained with Hematoxylin for 30 seconds, then bluing solution (NH_3_OH + MilliQ H2O, 2 mins) and Eosin for 1min followed by dehydration and mounting (Permount). Visualization of mammary gland ducts was performed using the Leica Widefield microscope, and ductal quantification was conducted using Fiji.

### Mammary gland whole mounts and branching analysis

Inguinal mammary glands were removed, adhered to a Superfrost Plus Microscope slide (Fisherbrand Cat. No. 1255015) and incubated in Carnoy’s Fixative overnight at room temperature. Mammary glands were then incubated in 70% ethanol, 50% ethanol, then DI water for 15 minutes each. Fixed mammary glands were stained overnight with Carmine Alum at room temperature, then dehydrated in 15-minute incubations in increasing ethanol grades, followed by dehydration in xylene and mounted in Permount.

Images of the entire mammary gland wholemount for both parous and nulliparous samples were imported as a TIFF file onto the Fiji/ImageJ software, where a three-by-three tiled ROI was chosen from a consistent distance from the lymph node. The ROI was processed to remove background noise. The image was skeletonized, and the branches dilated to improve clarity. A Sholl analysis (neuroanatomy) plugin was used on the skeletonized image, and a vertical radius from the side of the image was set as a consistent measurement throughout all samples to account for decreased branching normally seen at the edges of an image. After setting the starting radius as zero microns and selecting the best-fitting Sholl methods, the Sholl analysis program was run. This generated a “Sholl Log Plot,” which indicated branching intersections over an area, as well as a “Sholl Regression Coefficient (k)”, which described branching complexity. A k value closer to zero indicated a higher branching complexity. A “Sholl Mask (heat map)” was also generated to visually depict regions of higher branching complexity, where the red areas indicated maximal branching.

### Immunofluorescent staining of mammary gland paraffin sections

Mammary glands were fixed in 4% paraformaldehyde (PFA) in PBS and embedded in paraffin for immunostaining. 5μm sections were deparaffinized, dehydrated, and autoclaved for 15 min in Tris-EDTA buffer [10mM Tris 1mM EDTA (pH 9.0)] for antigen retrieval. Tissue sections were incubated overnight at 4°C with primary antibodies diluted in tris-buffered saline (TBS) + 5% bovine serum albumin (BSA) (antibodies are listed in table 5). Samples were subsequently washed (2X) with TBS + 0.05% Tween for 10 min and were incubated with donkey anti-rat Alexa Flour 647 (1:400), donkey anti-chicken Alexa Flour 488 (1:400), donkey anti-rabbit Alexa Flour 594 (1:400) conjugated secondary antibodies (Jackson ImmunoResearch Laboratories) in TBS + 5% BSA 0.1% Tween for 1 hour at room temperature (RT). Samples were subsequently washed (3X) with TBS + 0.1% Tween and were incubated with DAPI in TBS + 5% BSA + 0.1% Tween (1:10,000) for 10 min. All the immunofluorescence sections and cells were mounted in Fluoromount-G (Genesee). Images were acquired by a Solamere Spinning Disk Confocal microscope. Images were processed using Fiji.

### Immunofluorescent staining of primary MEC organoids

Organoids were collected at day 12-14 in culture and recovered from the growth factor-reduced Matrigel by incubating with Cell Recovery Solution (Corning) for 1 hour on ice with gentle shaking. 5mL collection tubes and pipette tips were precoated in PBS + 5% BSA. Organoids were washed with cold PBS (2X) before being fixed with 4% PFA in PBS for 45 min on ice. Fixed organoids were then washed in PBS (1X) and incubated with 0.2% glycine in PBS for 20 min at room temperature. After a final wash, fixed organoids were resuspended in 0.05% sodium azide in PBS and stored at 4°C.

Organoids were permeabilized in cold 100% methanol for 10 minutes on ice, washed PBS (1X), and incubated in blocking buffer (0.1% BSA, 0.3% Triton-X, 5% normal donkey serum) for 3h at room temperature with light shaking followed by overnight incubation with primary antibodies in blocking buffer at 4°C with light shaking. The next day they were washed (3X) in PBS + 0.3% Triton-X and incubated overnight with secondary antibodies in a blocking buffer at 4°C with light shaking in the dark. Organoids were again washed (3X) in PBS + 0.3% Triton-X and incubated with DAPI for 10 min at room temperature. After a final round of washes (3X), organoids were transferred to µ-Slide 8 Well Glass Bottom slide (Ibidi) pre-coated in poly-L-lysine (Sigma-Aldrich) and imaged on a Solamere Spinning Disk Confocal microscope. Images were processed using Fiji.

### Organoid assays

Primary MECs were sorted into complete organoid media as previous described^68,69^ (Advanced DMEM F/12, 10% FBS, 1% PSA, 50ng/mL EGF, 100ng/mL Noggin, 250ng/mL R-Spondin-1, 1X N2, 1X B27, 1X GlutaMAX, 10mM HEPES) supplemented with Y-27632 (10uM). 1,000 cells were plated in 96-well ultra-low attachment plates seeded with a 50uL mixture of growth factor-reduced Matrigel and irradiated L-Wnt3a-secreting-3T3 feeder cells (11,000 feeder cells per 50uL growth factor-reduced Matrigel). Organoids were cultured at 37C, 5% CO2 with added humidity. On day 4, Y-27632 supplemented media was removed and replaced with complete organoid media. Fresh media was added every other day thereafter. Basal and luminal organoids were imaged at day 6 and 10 in culture, respectively, on a Zeiss Live Cell microscope and analyzed on Biodoc.AI to collect data on organoid count and average size. For IL33 treatment experiments, organoids were cultured as described above with the addition of IL33 (0ng/mL, 5ng/mL, 10ng/mL).

### Mammary gland digestion and processing

L2-5 and R2-5 mammary glands were harvested, minced, and chemically digested overnight in Advanced DMEM F/12 with 1% PSA, gentle collagenase/hyaluronidase, and DNAse I at 37C, 5% CO2, and added humidity as previously described^70^. Briefly, partially digested glands were then mechanically digested by pipetting with a serological pipette until no tissue pieces were visible. Digested glands were washed with staining buffer (Hank’s Balanced Salt Solution, 2% Bovine Calf Serum, 1% PSA) and centrifuged (1500RPM) at 4C for 5 minutes. Red blood cells were lysed with 5mL of ACK Lysis buffer for 5 minutes, and cells were washed with 15mL of staining buffer. Cells were treated with 0.25% Trypsin with EDTA and gently pipetted continuously for 2-3 minutes to digest the basement membrane. Cells were then treated with DNAse I and Dispase and pipetted continuously for 2-3 minutes to prevent clumping. The single-cell suspension was then filtered through a 40um mesh strainer and pelleted via centrifugation (1500RPM, 4C, 5 minutes). Cells were then resuspended in staining buffer and transferred to FACS tubes for staining.

### Fluorescence-activated cell sorting (FACS)

Cells were stained with antibodies listed in Supplemental Table 5 for 15 minutes at room temperature, as previously described^68,69^. Stained cells were then washed with a staining buffer, resuspended with DAPI (1:10,000), filtered, and analyzed on a BD Biosciences FACSAria cell sorter. See Extended Data Fig. 2 for FACS gating strategies. Data were analyzed using FlowJo software (10.10.0).

### Lentiviral Infection

For *Trp53* knockdown experiments, pSicoR-GFP-sh*Trp53* was a gift from Tyler Jacks (Addgene plasmid #12090; http://n2t.net/addgene:12090; RRID:Addgene_12090). Viruses were produced in 293T cells using the second-generation lentiviral system and transfection using Lipofectamine 2000 (Life Technologies) as previously described^70^. Supernatants were collected at 48 hours, filtered with a 0.45μm filter, and precipitated with lentivirus precipitation solution (Alstem LLC) per the manufacturer’s instructions. Viral titers were determined by flow cytometry analyses of 293T cells infected with serial dilutions of concentrated virus.

### Single-cell RNA-seq cell preparation and sequencing

Libraries were generated using the 10x Genomics Chromium Next GEM Single Cell 5’ Reagent Kits v2 (Dual Index) (cat#1000264) following the manufacturer’s protocol. Single-cell suspensions were prepared from mammary glands as described above and encapsulated using the Chromium Controller to generate Gel Beads-in-Emulsion (GEMs), allowing for barcoding of individual cells. Post-GEM generation, reverse transcription (GEM-RT) was performed within each GEM, followed by cleanup and cDNA amplification. The cDNA underwent fragmentation, end repair, and A-tailing. Dual index adapters were ligated to the cDNA fragments, which were then amplified using PCR. The libraries were quantified using a Qubit Flex Fluorometer (Invitrogen Q33327) and their size distribution was assessed using an Agilent TapeStation 4200. Sequencing libraries were constructed to target 200 million paired-end reads (400 million total) to achieve a coverage of 20,000 reads per cell from a targeted capture of 10,000 cells. Sequencing was performed on an Illumina NovaSeq 6000 following the manufacturer’s recommendations.

### Single-cell data pre-processing and quality control

Sequencing data was processed using the 10x Genomics Cell Ranger pipeline. Reads were aligned to the mouse reference genome (mm10). We performed quality control for downstream analysis, removing (1) genes that were detected in less than 3 cells, (2) cells with less than 200 genes, (3) cells with gene counts < 600 or > 8,000, (4) cells with total counts of UMIs per cell < 2,000 or > 12,000, and (5) cells with mitochondrial gene ratio > 1.5%. The mitochondrial gene ratio is defined as the percentage of UMIs mapped to mitochondrial genes compared to non-mitochondrial genes within each cell. Doublets were identified using Scrublet^73^, resulting in the removal of 588 cells. Data preprocessing and analysis steps below were implemented using the Scanpy framework version 1.19^71^.

### Single cell data integration, dimension reduction, and cell type annotation

We combined our single cell data set with an 18 month nulliparous pre-processed single cell data set from Tabula Muris Senis^34^. The UMI counts for each cell were normalized using a target sum of 1e4, and log transformed, with an added pseudocount of 1. This resulted in a combined dataset of 10,001 cells and 13,892 genes. Principal component analysis (PCA) was conducted to produce a reduced dataset that was used as input to correct for technical variation due to samples using Harmonypy, version 0.0.4^72^. Neighborhood graphs were calculated on batch corrected reduced data and used to conduct Leiden^73^ unsupervised clustering, with a resolution of 0.09, resulting in the identification of nine distinct clusters. These clusters were manually annotated using well-established marker genes. Data was visualized using Uniform Manifold Approximation and Projection (UMAP, version 0.5.2)^74^.

### Cell-cycle analysis, basal and luminal gene scoring

Cell cycle enrichment analysis was conducted using the Regev lab cell cycle gene list^75^ to classify each cell into its corresponding cell cycle phase. To classify epithelial cells, we used previously published marker genes identified for Basal, HR-high Luminal, and HR-low Luminal cells from transcriptomic analyses of adult mammary epithelial cells (MECs)^35^. Enrichment scores for these marker genes were calculated for each of the four epithelial cell types (Basal epithelial, HR-high Luminal, HR-low Luminal, and Hybrid MECs). Cell cycle and epithelial cell type classifications were conducted using the ‘sc.tl.score_genes’ function from the Scanpy library^71^.

### Pseudotime inference and cellular potency prediction

Pseudotime analysis was performed on the epithelial cell types (basal, HR-low, HR-high, and hybrid). Partition-based graph abstraction (PAGA, version 1.2)^36^ analysis was performed using Scanpy (version 1.9.6). The diffusion map was computed using the basal cell type as the root cell. To ensure robustness, pseudotime trajectory was also inferred using Python’s implementation of Slingshot (pyslingshot, version 0.1.3)^37^, with basal cells as the start node. Slingshot pseudotime ordering scores were scaled between 0 and 1 for comparison with PAGA results. To characterize cellular potency, we used Python’s implementation of CytoTRACE 2 (version 1.0.0)^38^. Figures were generated using a combination of Scanpy^71^, Seaborn^76^, and matplotlib python packages^77^. To assess the statistical significance of the differences in pseudotime between epithelial cell types, we performed pairwise comparison using the Wilcoxon Rank Sum Test using Benjamini-Hochberg correction method to adjust the p-values and control the false discovery rate. For this analysis we used ‘scipy.stats.mannwhitneyu’ function^78^ for the Wilcoxon Rank Sum Test and ‘multipletests’ from the ‘statsmodels.stats.multitest’ library for Benjamini-Hochberg testing correction^79^.

### Library Construction, Quality Control and Bulk RNA Sequencing

Bulk RNA sequencing was performed by Novogene on sorted basal (CD49f^hi^/EPCAM^low-med^) and luminal (CD49f^low^/EPCAM^hi^) populations from 3m NP, 18m NP and 18m P mice (n=3 mice/group). RNA was isolated according to manufacturer’s instructions (Qiagen RNEasy Plus Micro Kit, Cat. No. 74034). Messenger RNA was purified from total RNA using poly-T oligo-attached magnetic beads. After fragmentation, the first strand cDNA was synthesized using random hexamer primers, followed by the second strand cDNA synthesis using either dUTP for directional library or dTTP for non-directional library^80^. For the non-directional library, it was ready after end repair, A-tailing, adapter ligation, size selection, amplification, and purification. For the directional library, it was ready after end repair, A-tailing, adapter ligation, size selection, USER enzyme digestion, amplification, and purification. The library was checked with Qubit and real-time PCR for quantification and bioanalyzer for size distribution detection. Quantified libraries will be pooled and sequenced on Illumina platforms, according to effective library concentration and data amount.

### Data Quality Control

Raw data (raw reads) of fastq format were processed through fastp software. In this step, clean data (clean reads) were obtained by removing reads containing adapter, reads containing ploy-N and low quality reads from raw data. Q20, Q30 and GC content were calculated. All the downstream analyses were based on the clean data with high quality.

### Reads mapping to the reference genome

Reference genome (GRCm39/mm39) and gene model annotation files were downloaded from genome website directly. Index of the reference genome was built usingHisat2 v2.0.5 and paired-end clean 1 reads were aligned to the reference genome using Hisat2 v2.0.5. We selected Hisat2^81^ as the mapping tool for that Hisat2 can generate a database of splice junctions based on the gene model annotation file and thus a better mapping result than other non-splice mapping tools.

### Quantification of gene expression level

featureCounts^82^ v1.5.0-p3 was used to count the reads numbers mapped to each gene. Gene expression was then converted to Fragments Per Kilobase of transcript sequence per Millions base pairs sequenced (FPKM), which takes the effects into consideration of both sequencing depth and gene length on counting of fragments. Analysis was performed by Novogene.

### Differential expression analysis

Differential expression^83^ analysis of two conditions (three biological replicates per condition) was performed using the DESeq2Rpackage^84^ (1.20.0) on raw gene expression values by Novogene. DESeq2 provides statistical routines for determining differential expression in digital gene expression data using a model based on the negative binomial distribution. Differentially expressed genes were identified by comparing 3m NP vs. 18m NP and 18m P vs 18m NP for both luminal and basal cell samples. The resulting P-values were adjusted using the Benjamini-Hochberg’s approach for controlling the false discovery rate. Genes with an adj-p<=0.1 found by DESeq2 were assigned as differentially expressed. Principal Component Analysis (PCA) plots and heatmaps of differentially expressed genes were generated using the NovoMagic analysis platform provided by Novogene. PCA plots are based on FPKM normalized gene expression values. Heatmaps of differentially expressed genes display log_2_(FC+1) values. Representative genes from the top 30 DEGs are shown. See Data tables 1-4 for complete gene list.

### Enrichment analysis of differentially expressed genes

Gene Ontology^85^ (GO) enrichment analysis of differentially expressed genes was implemented by the cluster Profiler R package, in which gene length bias was corrected. GO terms with corrected p-value less than 0.1 were considered significantly enriched by differential expressed genes.

### Statistical analysis

All graphs display the average as central values, and error bars indicate ± SD unless otherwise indicated. P values are calculated using paired or unpaired t test, ANOVA, Wilcoxon rank-sum test, and Mann-Whitney U test, as indicated in the figure legends. All P and Q values were calculated using Prism (10.2.2) or Python, unless otherwise stated. For animal studies, sample size was not predetermined to ensure adequate power to detect a prespecified effect size, no animals were excluded from analyses, experiments were not randomized, and investigators were not blinded to group allocation during experiments.

## DATA AVAILABILITY

Data generated or analyzed during this study are included in this published article (and its supplemental information files). Data needed to evaluate the conclusions in the paper are present in the paper and/or the Supplemental Materials. Source data files will be made available upon request. Single-cell and bulk RNA-sequencing data generated in this study have been deposited in the Gene Expression Omnibus with the primary accession code GSE272933 (bulk) and GSE272932 (single-cell). All bioinformatics tools used in this study are published and publicly available.

## ACKNOWLEDGMENTS

We thank Bari Nazario for her help in flow cytometry. The FACS Aria instrument was funded by NIH grant S10-1S10RR02933801. We thank Benjamin Abrams, UCSC Life Sciences Microscopy Center, RRID: SCR_021135 for technical support during image acquisition and processing. We also thank the animal facility core members for animal maintenance. We thank Camilla Forsberg, Lindsay Hinck, Aaron Newman and members of the Sikandar lab for critical feedback on the manuscript. The authors declare no competing interests. This work was supported by the Hellman Fellows Award and startup funds (to S.S.S), NIH T-32 (5T32GM133391-04) (to A.O), the Eugene Cota-Robles Fellowship (to C.M.R) and startup funds (to V.D.J). S.S.S is also supported by the NIH/NCI (R37CA269754).

## AUTHOR CONTRIBUTIONS

S.S.S. and A.O conceived and designed the study. A.O performed most of the experiments and analyzed the data with assistance from V.H.A and S.K and under the supervision of S.S.S. V.H.A collected samples for single-cell RNA sequencing and processed samples for bulk-RNA sequencing. P.M verified the differential gene expression analysis from bulk RNA sequencing data. C.M.R analyzed single-cell RNA sequencing data under supervision of V.D.J. A.O and S.S.S. wrote the manuscript with contributions from C.M.R and V.D.J. All authors commented on the manuscript.

## COMPETING INTERESTING

The authors declare no competing interests.

## EXTENDED DATA

Extended data figures 1-6

Tables 1-5

## Notes

### Competing Interest Statement

The authors have declared no competing interest.

