## Supplementary Figures for "Pregnancy Reduces Il33+ Hybrid Progenitor Accumulation in the Aged Mammary Gland"

**This PDF includes:**

Extended data figures 1-6

Table 5

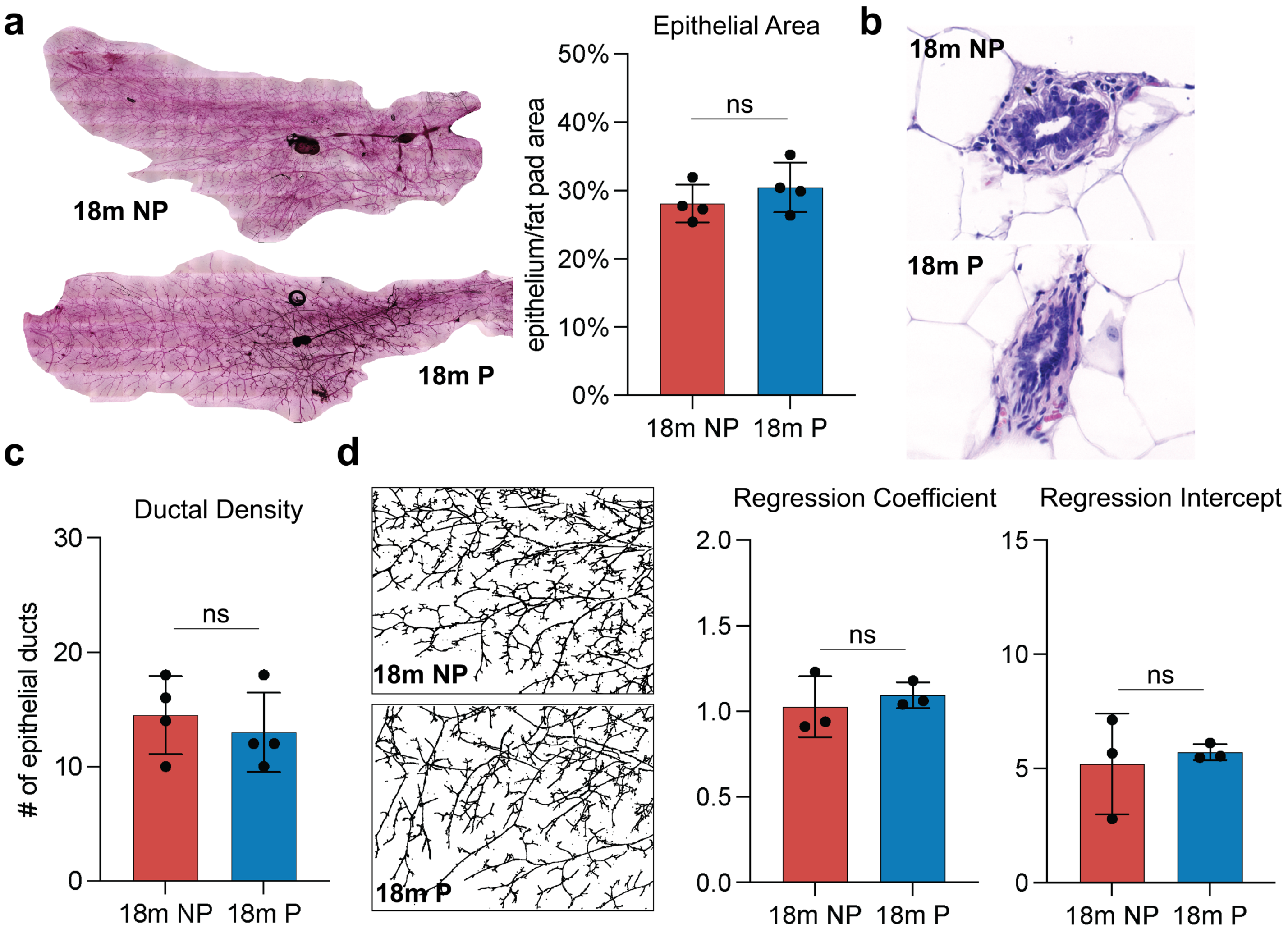


**Extended Data Figure 1**. **Wholemount and H&E analysis of aged, parous and nulliparous mammary glands.** (a) Representative wholemount images of cleared mammary glands from aged, parous and nulliparous mice stained with Carmine Alum (left) and the quantification of epithelial area from wholemount images (right). (b) Representative H&E images of epithelial ducts. (c) Quantification of ductal density per field of view from H&E images. (d) Representative images of skeletonized epithelial structures (left) used in Sholl analysis to quantify branching complexity (regression coefficient and regression intercept). Statistical significance was determined by performing unpaired t tests. * $p < 0.05$, ** $p < 0.01$, *** $p < 0.001$. n = 4 mice (a-c) or 3 mice (d). Scale bar = 1000 μm (a, d) and 100 μm (c).

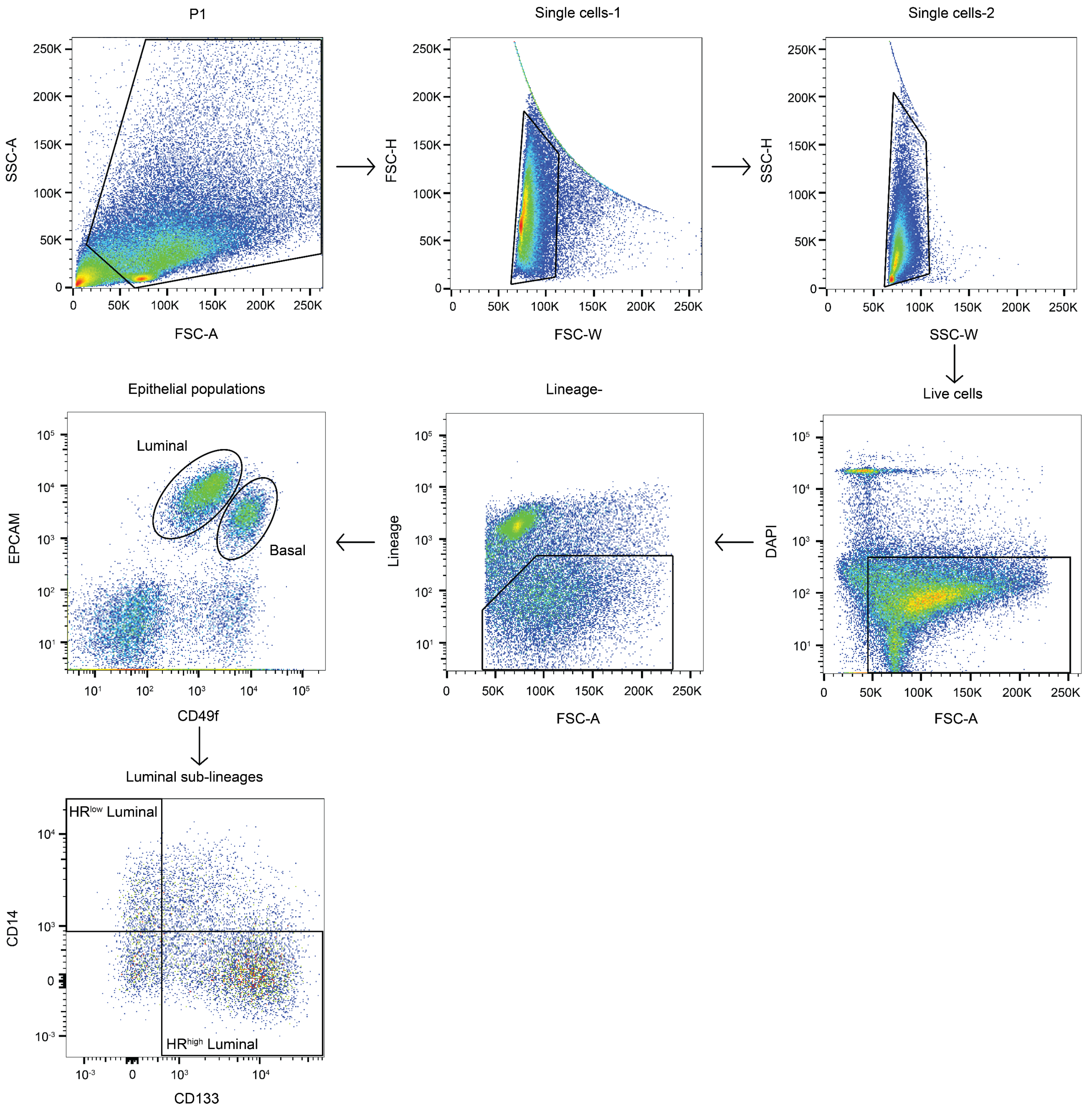


**Extended Data Figure 2. Gating strategy for flow cytometry and fluorescence-activated cell sorting experiments.** Single cell suspensions of mammary glands were analyzed on a BD Aria Cell Sorter. Cellular debris was removed in the “P1” SSC-A x FSC-A gate (top left). Single cells are isolated through two singlet gates using FSC-H x FSC-W and SSC-H x SSC-W (top row, middle and right panel). Live cells (DAPI-) are isolated (second row, right panel) and immune cells (CD45-Pacific Blue), red blood cells (Ter119-Pacific Blue), and endothelial cells (CD31-Pacific Blue) are gated our through a lineage(-) gate (second row, middle panel).

Basal (CD49f$^{hi}$/EPCAM$^{med/low}$) and luminal (CD49f$^{low}$/EPCAM$^{hi}$) epithelial cells are then isolated from the lineage(-) population (second row, left panel). Luminal cells are divided into HR$^{high}$ and HR$^{low}$ sublineages using CD14-APC/Cy7 and CD133-PE, respectively (third row, left panel). HR$^{high}$ and HR$^{low}$ luminal gates were set based on FMO negative controls.

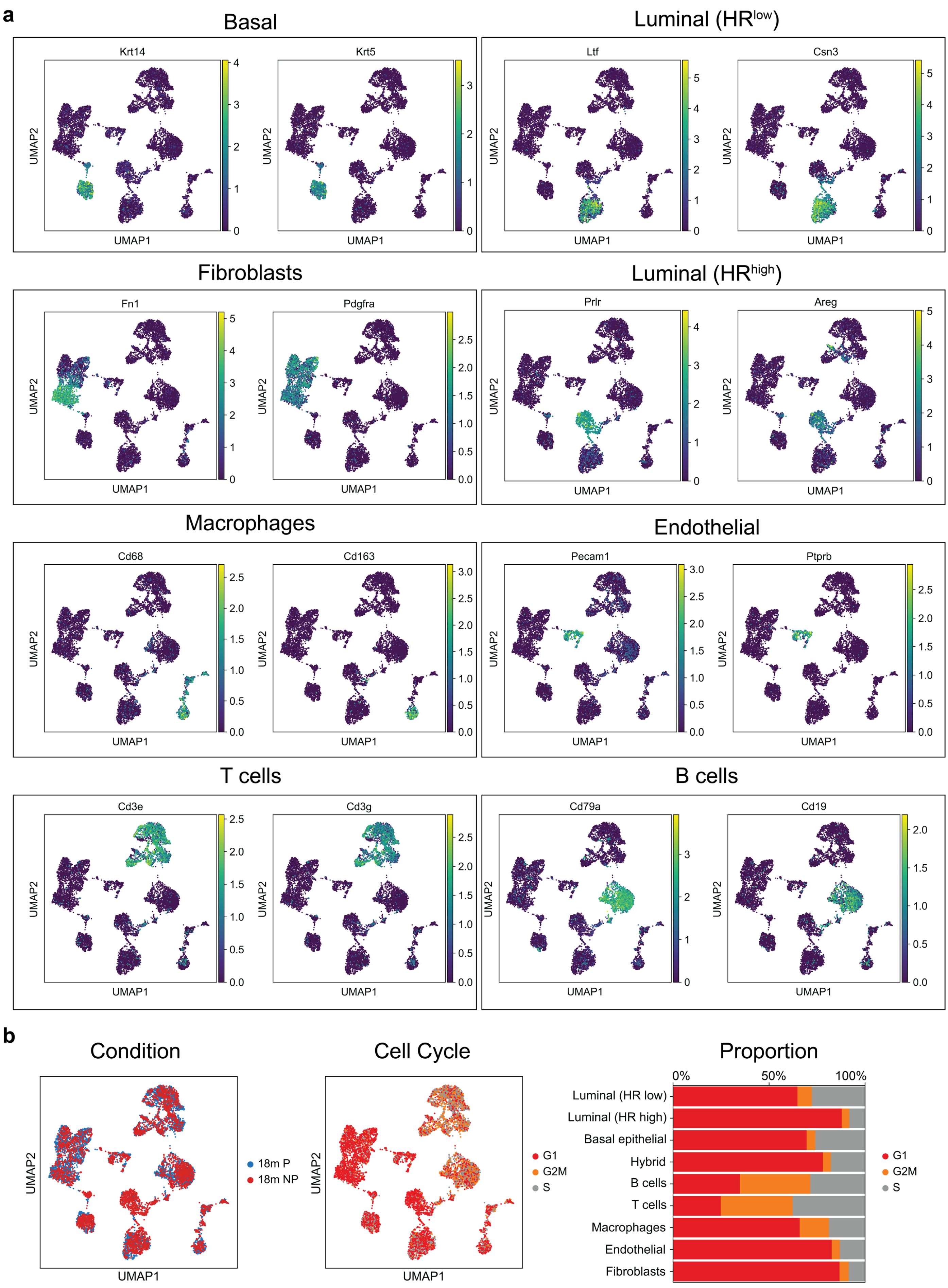

a
Basal
Krt14
Krt5
Luminal (HRlow)
Ltf
Csn3
Fibroblasts
Fn1
Pdgfra
Luminal (HRhigh)
Prlr
Areg
Macrophages
Cd68
Cd163
Endothelial
Pecam1
Ptprb
T cells
Cd3e
Cd3g
B cells
Cd79a
Cd19
UMAP1
UMAP2
b
Condition
18m P
18m NP
Cell Cycle
G1
G2M
S
Proportion
0%
50%
100%
Luminal (HR low)
Luminal (HR high)
Basal epithelial
Hybrid
B cells
T cells
Macrophages
Endothelial
Fibroblasts

**Extended Data Figure 3. Cell type annotation and cell cycle analysis of single cell RNA sequencing data.** (a) UMAP plots colored by expression of lineage markers used for cell type annotation (Basal, Luminal-HR$^{low}$, Luminal-HR$^{high}$, Fibroblasts, Macrophages, Endothelial, T cells, and B cells). (b) UMAP plot colored by condition (18m P and 18m NP, left) and cell cycle stage (middle). The proportion of cells in G1, G2M, and S phase are represented in a bar chart and stratified by cell type (right).

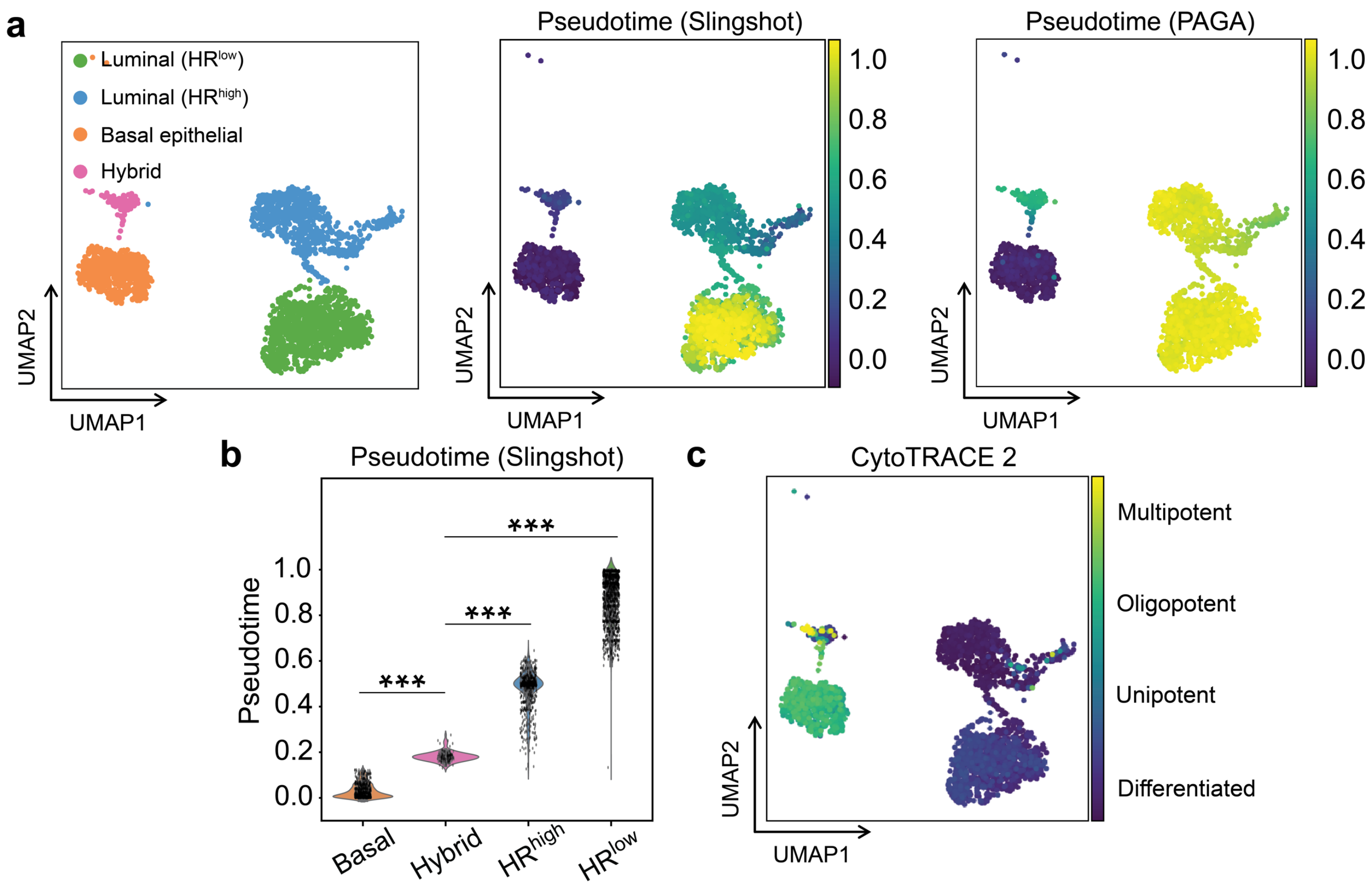


**Extended Data Figure 4. Pseudotime analysis and CytoTRACE 2 analysis of MECs.** (a) UMAP plot of epithelial cells colored by cell type (left), Slingshot pseudotime score (middle), and PAGA pseudotime score (right). (b) Violin plots of Slingshot pseudotime scores across cell types. (c) UMAP plot of epithelial cells colored by CytoTRACE 2 potency score.

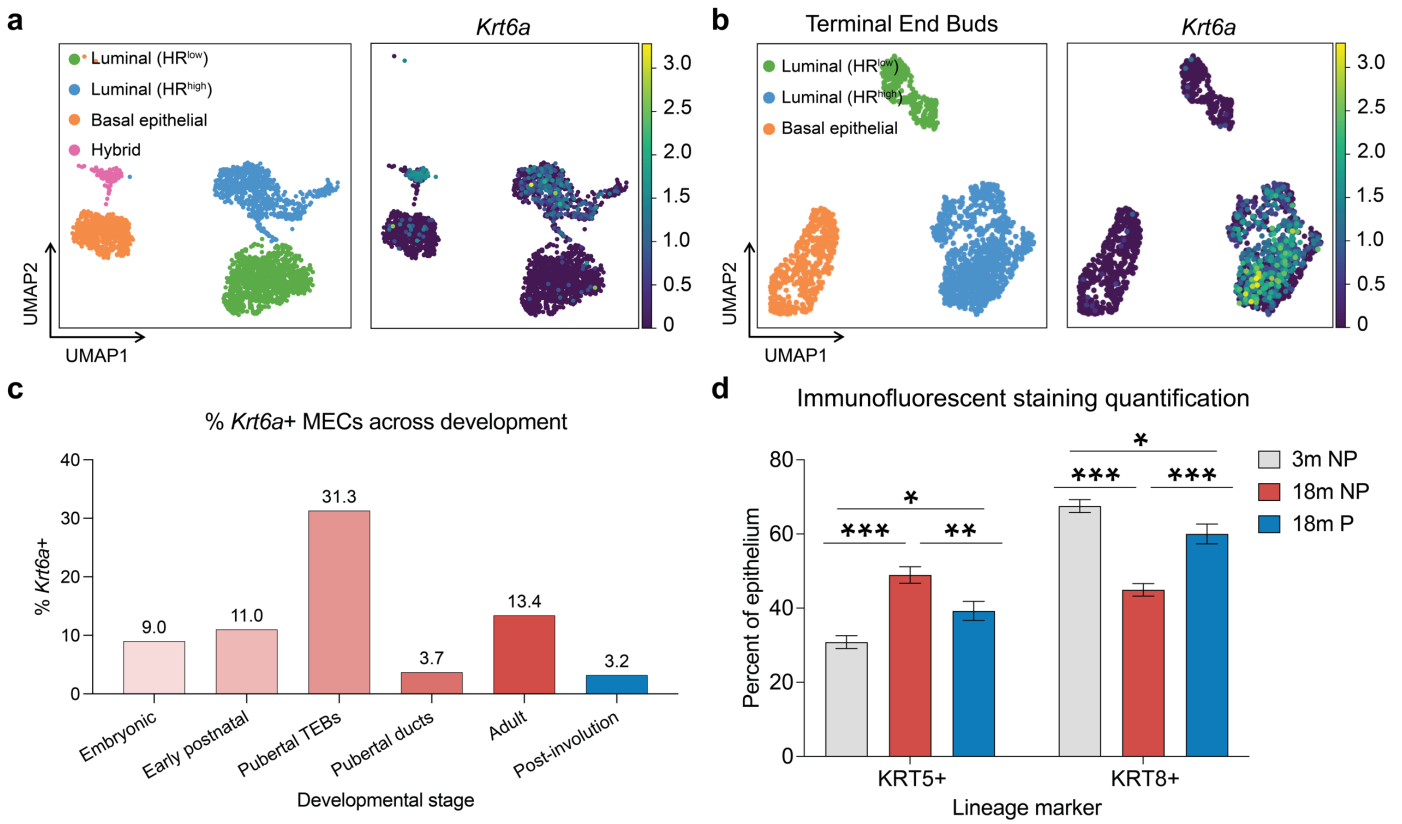


**Extended Data Figure 5. KRT6a, KRT5 and KRT8 at different developmental points in MECs.** (a) UMAP plot of epithelial cells from 18m NP and P mice, colored by cell type (left) and *Krt6a* expression (right). (b) UMAP plot of epithelial cells from TEBs colored by cell type (left) and *Krt6a* expression. (c) Quantification of *Krt6a*+ MECs across developmental stages (Pal et al., 2017 dataset). (d) Quantification of basal (KRT5+) and luminal (KRT8+) epithelial cells from immunofluorescent staining. Error bars represent +/- S.E.M. * p <0.05, ** p < 0.01, *** p < 0.001.

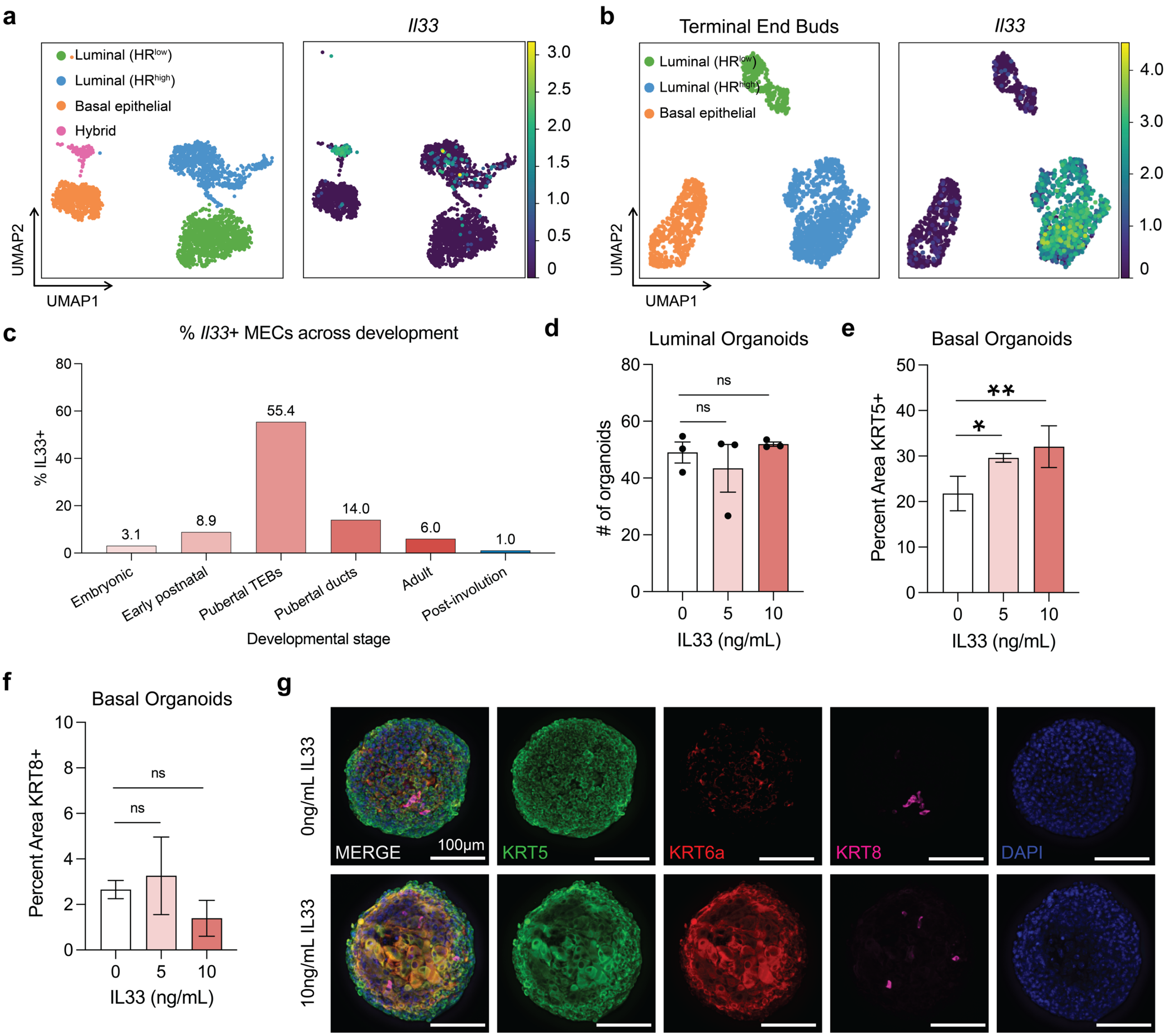


**Extended Data Figure 6. Expression of *Il33* at different developmental timepoints and quantification of KRT8 and KRT5 in organoids post IL33 treatment.** (a) UMAP plot of epithelial clusters colored by cell type (left) and *Il33* expression (right). (b) UMAP plot of epithelial cells from TEBs, colored by cell type (left) and *Il33* expression (right) (Pal et al., 2017 dataset). (c) Quantification of *Il33*+ MECs across developmental stages (Pal et al., 2017 dataset). (d) Quantification of organoid formation for primary luminal cells treated with IL33. (e) Quantification of percent KRT5+ area from IF experiments on IL33-treated basal organoids. (f) Quantification of percent KRT8+ area from IF experiments on IL33-treated basal organoids. (g) Representative IF images of IL33-treated basal organoids. Error bars represent +/- S.E.M. * $p < 0.05$, ** $p < 0.01$, *** $p < 0.001$.

**Table 5. Antibodies**

| Antibody | Dilution | Clone | Manufacturer | Catalog Number |
|---|---|---|---|---|
| PE/Cyanine7 anti-mouse CD326 (EPCAM) | 1:200 | G8.8 | Biolegend | 118216 |
| APC anti-human/mouse CD49f | 1:100 | GoH3 | Biolegend | 313616 |
| PE anti-mouse CD133 | 1:100 | 315-2C11 | Biolegend | 141203 |
| APC/Cyanine7 anti-mouse CD14 | 1:50 | Sa14-2 | Biolegend | 123318 |
| PE anti-mouse/rat CD61 Antibody | 1:100 | 2C9.G2 | Biolegend | 104308 |
| Pacific Blue anti-mouse TER-119/Erythroid | 1:200 | TER-119 | Biolegend | 116232 |
| Pacific Blue anti-mouse CD31 | 1:200 | 390 | Biolegend | 102422 |
| Pacific Blue anti-mouse CD45 | 1:200 | S18009F | Biolegend | 102422 |
| Rabbit anti-mouse KRT6a | 1:50 | - | Biolegend | 905701 |
| Chicken anti-mouse KRT5 | 1:100 | - | Biolegend | 905901 |
| Rat anti-mouse KRT8 | 1:200 | TROMA-1c | Developmental Studies Hybridoma Bank | NA |
| Chicken anti-mouse IL33 | 1:100 | - | Invitrogen | PIPA547007 |
| Donkey anti-goat Alexa Fluor 488 | 1:400 | - | Invitrogen | A11034 |
| Donkey anti-chicken Alexa Fluor 488 | 1:400 | - | Jackson ImmunoResearch Laboratories | 703-005-155 |
| Donkey anti-rabbit Alexa Fluor 594 | 1:400 | - | Jackson ImmunoResearch Laboratories | 711-585-152 |
| Donkey anti-rat Alexa Fluor 647 | 1:400 | - | Jackson ImmunoResearch Laboratories | 712-605-150 |
